# Levetiracetam Modulates Brain Metabolic Networks and Transcriptomic Signatures in the 5XFAD Mouse Model of Alzheimer’s disease

**DOI:** 10.1101/2023.11.10.566574

**Authors:** Charles P. Burton, Evgeny J. Chumin, Alyssa Y. Collins, Scott A. Persohn, Kristen D. Onos, Ravi S. Pandey, Sara K. Quinney, Paul R. Territo

## Abstract

**INTRODUCTION:** Subcritical epileptiform activity is associated with impaired cognitive function and is commonly seen in patients with Alzheimer’s disease (AD). The anti-convulsant, levetiracetam (LEV), is currently being evaluated in clinical trials for its ability to reduce epileptiform activity and improve cognitive function in AD. The purpose of the current study was to apply pharmacokinetics (PK), network analysis of medical imaging, gene transcriptomics, and PK/PD modeling to a cohort of amyloidogenic mice to establish how LEV restores or drives alterations in the brain networks of mice in a dose-dependent basis using the rigorous preclinical pipeline of the MODEL-AD Preclinical Testing Core.

**METHODS:** Chronic LEV was administered to 5XFAD mice of both sexes for 3 months based on allometrically scaled clinical dose levels from PK models. Data collection and analysis consisted of a multi-modal approach utilizing ^18^F-FDG PET/MRI imaging and analysis, transcriptomic analyses, and PK/PD modeling.

**RESULTS:** Pharmacokinetics of LEV showed a sex and dose dependence in C_max_, CL/F, and AUC_0-∞_, with simulations used to estimate dose regimens. Chronic dosing at 10, 30, and 56 mg/kg, showed ^18^F-FDG specific regional differences in brain uptake, and in whole brain covariance measures such as clustering coefficient, degree, network density, and connection strength (i.e. positive and negative). In addition, transcriptomic analysis via nanoString showed dose-dependent changes in gene expression in pathways consistent ^18^F-FDG uptake and network changes, and PK/PD modeling showed a concentration dependence for key genes, but not for network covariance modeling.

**DISCUSSION:** This study represents the first report detailing the relationships of metabolic covariance and transcriptomic network changes resulting from LEV administration in 5XFAD mice. Overall, our results highlight non-linear kinetics based on dose and sex, where gene expression analysis demonstrated LEV dose- and concentration-dependent changes, along with cerebral metabolism, and/or cerebral homeostatic mechanisms relevant to human AD, which aligned closely with network covariance analysis of ^18^F-FDG images. Collectively, this study show cases the value of a multimodal connectomic, transcriptomic, and pharmacokinetic approach to further investigate dose dependent relationships in preclinical studies, with translational value towards informing clinical study design.

## INTRODUCTION

Subclinical epileptiform activity is commonplace in both mouse models of Alzheimer’s Disease (AD) and in the clinic (Minkeviciene et al., 2009;Vossel et al., 2016). Recent work has shown that sustained expression of the calcium binding protein, calbindin-D_28K_, buffers cytosolic calcium in mouse models of AD, while healthy controls showed decreased expression with age in subicular dendrites, thus forming a molecular basis of epileptiform activity in models of AD (Stafstrom, 2005;Angulo et al., 2020). Additional work has reported spike wave discharge activity in the 5XFAD mouse of model of AD, which is consistent with human AD epileptiform activity, where hippocampal hyper-excitability has been observed in human *APOE^E4/E4^*carriers (Tran et al., 2017). Importantly, this activity has been shown to localize to β-amyloid (Aβ) plaques, where neuronal dysregulation is thought to result from impaired synaptic inhibition, thus driving aberrant excitatory activity. Neurons distal to the plaques show abnormally low spike wave discharge activity (Palop et al., 2007), suggesting that Aβ plaque load may play an important role in epileptiform formation. Provided this, anti-epileptic drugs have been considered as a possible treatment paradigm and are hypothesized to attenuate synchronized hyperactivity in patients with AD (Palop, 2009). One drug in particular, Levetiracetam (LEV), received FDA approval in 2000 for the treatment of seizures and epilepsy in adults and adolescents (Kumar et al., 2023). Recent clinical trials administering LEV in AD patients have found improved cognitive function in patients with epileptiform activity (Vossel et al., 2021). In humanized amyloid precursor protein (*hAPP)* mice, LEV significantly reduced epileptiform activity in a dose-dependent manner and halted disease progression (Sanchez et al., 2012). Combined, these data suggest that LEV treatment may be efficacious in attenuating disease progression in 5XFAD mice that overexpress Aβ and suggest that prophylactic treatment with LEV may provide experimental evidence as a treatment for late onset AD (LOAD) [6].

Clinically, hyper-excitability has been associated with seizures in AD patients (Lippa et al., 2010;Vossel et al., 2016;Toniolo et al., 2020). This phenomenon is also observed in AD mouse models such as 5XFAD and *APP/PS1* (Schneider et al., 2014;Klee et al., 2020). LEV’s acute administration in amyloid mouse models has shown attenuation of abnormal spike wave activity (Klee et al., 2020), while chronic treatment via continuous mini-pump infusion has been associated with the reversal of synaptic loss and behavioral impairment in *hAPP* mice (Sanchez et al., 2012). However, prior preclinical studies have not fully characterized the pharmacokinetics (PK) of LEV and its impact on gene transcriptomic and brain metabolic network alterations that are crucial for predicting the observed nonlinear dose efficacy in clinical studies. Notably, single-daily dosing of LEV may not maintain exposure levels, and could result in elevated C_min_/C_max_ ratios, thus reducing drug effectiveness and potentially resulting in C_max_ dependent toxicities.

Although the precise mechanism of action of LEV in AD is not fully understood, it is known to modulate the release of synaptic neurotransmitters by antagonizing SV2A receptors (Lynch et al., 2004;Kumar et al., 2023). Moreover, neuromodulation is thought to occur through the inhibition of presynaptic calcium channels, reducing impulse conductivity and thus enhancing the preferential transmission of low-frequency signals (García-Pérez et al., 2015). During the repurposing of LEV for use in mild cognitive impairment (MCI) studies, a nonlinear dose-dependent response curve was observed in the form of an inverted paraboloid (Bakker et al., 2012;Bakker et al., 2015;Magalhães et al., 2015;Musaeus et al., 2017). A study with control and MCI participants imaged with functional magnetic resonance imaging (MRI) while performing a memory task revealed that the low (62.5 mg twice daily (BID)) and moderate (125 mg BID) doses of LEV restored normal hippocampal activity in the dentate gyrus and CA3 regions. However, this effectiveness was not observed in the highest (250 mg BID) dose studied, suggesting a resurgence in nonequilibrium synchronization in the hippocampus (Bakker et al., 2012).

Previous work from our lab has explored the pharmacokinetic (PK) and pharmacodynamic (PD) relationships of LEV in a mouse model of AD that could be used to better predict clinical efficacy (Onos et al., 2022). We evaluated LEV using the validated pipeline of the Model Organism Development and Evaluation for Late Onset Alzheimer’s Disease (MODEL-AD) Preclinical Testing Core (PTC), where we determined the PK of LEV in the 5XFAD mouse model, permitting the determination of dose, frequency, and duration for chronic studies. We performed PD assessment followed chronic administration, where ^18^F-AV45 and ^18^F-FDG PET/MRI showed dose-dependent reduction in amyloid deposition and glucose uptake across imaging cohorts. Additional sex and dose dependencies were seen in nanoString estimates of transcriptional changes, which underpin the PD changes. Though these results showed significant alterations in PK and PD as a function of treatment, methods to elucidate how LEV modified gene and associated brain networks and how this related to blood and brain drug concentrations (i.e. PK/PD) was not investigated (Onos et al., 2022).

Recent work from our group has applied a metabolic network covariance modeling approach in the 5XFAD model across lifespan (i.e. 4, 6, and 12 mos) (Chumin et al., 2023). This work permits the interrogation of network changes at the whole brain and sub-network levels, allowing greater insights into the network changes that occur with disease progression. Here, we applied this method to investigate LEV treatment and subsequent interregional metabolic changes in 5XFAD mice of both sexes. Our goal was to establish a clear dose/concentration-dependent relationship of LEV treatment in conjunction with alterations in brain metabolic uptake network and gene expression to better predict clinical efficacy, which is not currently possible using traditional analytical approaches.

## METHODS

All studies adhered to the ARRIVE guidelines and received approval from the Institutional Animal Use and Care Committees (IACUC at their respective locations).

### Housing conditions and cohort generation at Indiana University (IU)

Adult male 5XFAD mice, female 5XFAD mice, and non-transgenic WT controls were produced through a breeding program at Indiana University (IU). This involved mating male 5XFAD mice (JAX MMRRC stock #:34848) with female C57BL6/J mice (JAX MMRRC stock #:000664). The animals were accommodated in cages, with up to five mice per cage, and were provided with SaniChip bedding. Throughout the dosing studies, the mice continued to reside in group housing arrangements. The colony room maintained a 12:12 Light:Dark schedule, with lights turning on at 6:00 am. One cohort was studied, designed for the endpoint of ^18^F-FDG PET.

### Housing conditions and cohort generation at The Jackson Laboratory (JAX)

Adult male 5XFAD mice, female 5XFAD mice, and non-transgenic WT controls were bred at JAX using the same breeding methods as those employed at IU. Initially, mice were accommodated in duplex cages with pine bedding, allowing up to five mice per side. The colony room adhered to a 12:12 Light:Dark schedule. For chronic dosing studies, each treatment group consisted of approximately 10-15 mice per sex. To ensure randomization, both treatment and sex were randomized across two cohorts, which were staggered by a 4-week interval. Each cohort was comprised of 5-8 mice per sex per treatment. One week prior to the beginning of the study, the mice were individually housed and subsequently transported to a colony room located adjacent to the behavioral testing facility.

### Levetiracetam Pharmacokinetic Studies

*In vivo* PK sampling for LEV was initially carried out at JAX following dosing and serial sampling in 6-month-old male and female 5XFAD mice. LEV (Sigma # L8668-100mg; Lot # 051M4742V) was dissolved in sterile saline (vehicle). Mice (with n=3 dose per sex) were administered doses of 10, 30, and 100 mg/kg (dose volume 10 mL/kg) via oral gavage as described previously (Onos et al., 2022). Serial plasma samples were collected via the tail vein prior to dosing and at 0.25, 0.5, 1, 2, 4, 6, and 24 hours post-dosing. Mice were euthanized at 24 hours, brains excised, frozen using dry ice, and stored at −80°C. Samples were shipped to IU for quantification of LEV and PK analysis per our previous work (Onos et al., 2022).

### Levetiracetam Quantification

LEV and etiracetam (ECA) concentrations were determined in plasma and brain samples using LC/MS/MS, with temazepam as an internal standard as described previously (Onos et al., 2022). Standard curves were established over a concentration range of 0.3-30000 ng/mL for plasma samples and 0.8-800 ng/g for brain homogenate samples. The inter-day precision ranged from 5.3% to 15.4% for LEV and 10.7% to 17.0% for ECA, while the inter-day accuracy ranged from 88.1% to 108.0% for LEV and 88.4% to 103.0% for ECA.

### Pharmacokinetic Modeling

PK parameters were initially calculated using standard noncompartmental analysis (NCA) performed with WinNonlin (Phoenix 64, build 8.0.0.3176), as we have previously described (Onos et al., 2022). To estimate chronic exposure, a population pharmacokinetic analysis was conducted in Monolix 2023R1 (Lixoft). Apparent clearance (CL/F) and volume (V/F) were allometrically scaled to weight with fixed coefficients of 0.75 and 1.0, respectively. Dose and sex were tested as covariates on CL/F and V/F. Model and covariate selection was based on reduction of objective function value of at least 3.84 (χ^2^, *p*<0.05), goodness of fit plots, covariate vs. eta plots, and visual predictive checks. Clearance for individual mice following chronic treatment with levetiracetam was calculated in Sumlx (2023R1, Lixoft) population estimates (thetas), and area under the concentration time (AUC) calculated as dose-normalized clearance. Data were analyzed and plotted in R version 4.3.0 (Team, 2023).

### Magnetic Resonance Imaging

High-contrast gray matter images were acquired 2 days prior to PET imaging, where mice were induced with 5% isoflurane in medical oxygen, placed on the head coil, and anesthesia was maintained with 1– 3% isoflurane for scan duration. High-resolution T2-weighted (T2W) MRI images were acquired using a 3T Siemens Prisma clinical MRI scanner outfitted with a dedicated 4-channel mouse head coil and bed system (RAPID MR, Columbus, OH, United States) per our previous work (Oblak et al., 2021;Onos et al., 2022).

### In vivo PET Imaging and Analysis

Regional brain glycolytic metabolism was monitored using 2-^18^F-fluoro-2-deoxy-d-glucose (^18^F-FDG), where clinical unit doses ranging from 185 MBq (5 mCi) were purchased from PETNet Indiana (PETNET Solutions Inc.). In all cases, mice were fasted for a minimum of 12 h prior to tracer administration. Mice were injected IP with 3.7–11.1 MBq (100–300 uCi) with ^18^F-FDG and allowed 30 min of uptake in an isothermal cage. For imaging, mice were then anesthetized with 5% isoflurane gas and maintained at with 1–3% isoflurane per our previous work (Oblak et al., 2021;Onos et al., 2022), and scanned on the IndyPET3 (Rouzes et al., 2004) scanner for 15 min. Images were calibrated, and decay- and scatter-corrected PET images were reconstructed into a single-static image volume according to our previous work (Oblak et al., 2021;Onos et al., 2022). All images were co-registered using a rigid-body mutual information-based normalized entropy algorithm with 9° of freedom and mapped to stereotactic mouse brain coordinates according to our previous work (Oblak et al., 2021;Onos et al., 2022). Post-registration, 56 regions were extracted via Paxinos and Franklin’s 2007 brain atlas and averaged to yield 28 bilateral regions (Paxinos and Keith B. J. Franklin, 2007). Standardized Uptake Value Ratios (SUVr; normalized to cerebellum) were computed for all regions relative to cerebellum, and principal component analysis was performed to provide data reduction for all PET regions. Consensus regions which explained 80% of the variance observed across all regions studied were selected for regional interrogation via MANOVA, correcting for multiple comparisons with a Bonferroni correction. To assess whole brain and sub-network changes in ^18^F-FDG images, metabolic covariance analysis was conducted according to our recent work and open source tools (Chumin et al., 2023). Pearson’s correlation between z-scores of region pairs was calculated for all pairwise interactions within each cohort to generate a covariance adjacency matrix (see Figures. 3, 5-7). To assess metabolic network characteristics, a correlation threshold of *p*<0.05 was applied to adjacency matrices, with only significant edges surviving (Figure 3). Multiple alpha values were tested, but thresholds more restrictive than *p*<0.05 yielded graphs too sparse for meaningful analysis. The nodal degree, positive strength, negative strength, and clustering coefficient was computed for every brain region and the global distribution of nodal characteristics were compared across sex and treatment using 2-sample Kolmogorov-Smirnov (KS) tests. To evaluate the sub-network modules, littermate control (WT) community partitions were generated via multi-resolution consensus clustering (MRCC) analysis (Jeub et al., 2018) and imposed on comparison groups. In sex comparisons, male littermate control partitions were used. Mean metabolic SUVr were compared between like communities via ANOVA, with a Bonferroni correction.

### Drug Administration for Chronic Studies

Mice were weighed daily and received oral gavage of LEV (SelleckChem # S1356, bulk lot # S135602) dissolved in sterile saline twice daily (BID) for 3 months. LEV was administered between 7:00 am – 9:00 am and 3:00 pm – 5:00 pm, with a dose volume of 10 mL/kg. In all cases, LEV was formulated in sterile saline weekly, and vials were blinded (i.e. A, B, C, and D (JAX) or Blue, Red, Yellow, and Green (IU)) in accordance with ARRIVE guidelines (Percie du Sert et al., 2020). Drug stability was determined by IU Clinical Pharmacology Analytical Core (CPAC) and was shown to be stable in the final formulation for a one-week period. Throughout the chronic dosing period, animals were closely monitored for potential signs of toxicity or drug-related side effects, where the attrition was very low (n = 3). This was not specific to a dose level and occurred either due to dermatitis or lung puncture during oral gavage.

### Terminal Tissue Collection

Upon study completion plasma and brain tissue samples were collected from the subjects under isoflurane anesthesia 30 min after the final LEV dose. Bioanalytical analysis was conducted via CPAC for terminal plasma and the right brain hemisphere to confirm PK data. For transcriptional profiling, homogenates from the left hemispheres were quantified using a customized nanoString nCounter^®^ Mouse AD panel designed to identify changes in gene expression associated with clinical LOAD. Differential gene expression was assessed with consideration of genotype, sex, and treatment per our previous work (Onos et al., 2022).

### nanoString Gene Expression Profiling and Analysis

Methods for this assay have been published previously (Onos et al., 2022). Data were analyzed with the use of QIAGEN IPA (QIAGEN Inc., https://digitalinsights.qiagen.com/IPA).

### Rigor and Reproducibility

During study execution and throughout data analysis, all technicians adhered to the ARRIVE guidelines (Percie du Sert et al., 2020) and were unaware of the genotype and drug dosage information.

## RESULTS

### Population Pharmacokinetics

LEV concentration-time data fit a 1-compartment first order absorption model (Figure 1). After accounting for weight, dose (but not sex) was found to be a significant covariate on oral clearance (CL/F) and apparent volume of distribution (V/F), as indicated by reduced objective function value and improved covariate vs. η plots. As shown in **Figure 1B**, the predicted AUCs were significantly correlated with 0.5 hour plasma concentrations following twice-daily dosing of levetiracetam in mice housed at JAX (p<0.001, R^2^=0.71)

**Figure 1.**
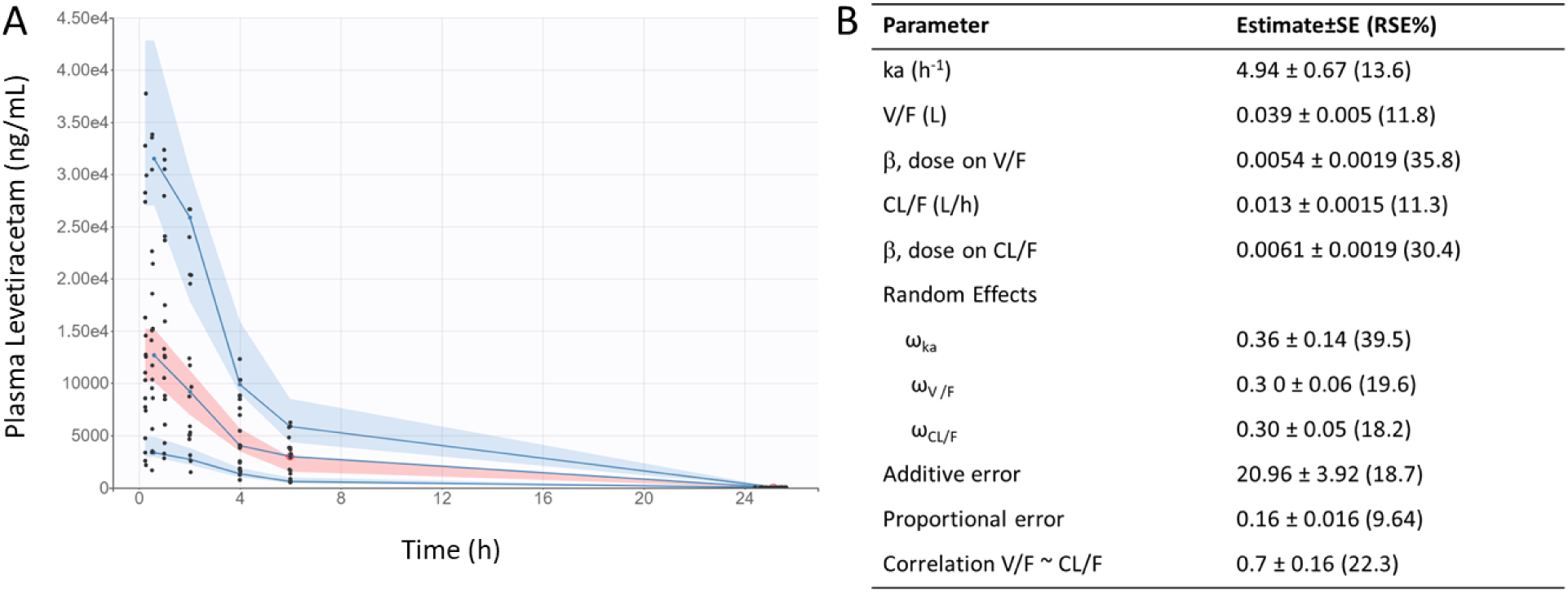
Levetiracetam pharmacokinetics: (A) Visual predictive check demonstrates reasonable fit of population pharmacokinetic model to data. Points indicate individual observed data, solid lines represent median, 5th, and 95th percentiles of observed data, pink shaded area indicates 90% prediction interval of the median, blue shaded areas indicated 90% prediction intervals of the 5th and 95th percentiles of predicted data. (B) Final pharmacokinetic parameter estimates for population model of levetiracetam.

### General Linear Modeling of Brain ^18^F-FDG PET

In chronic LEV-treated 5XFAD mice of both sexes, quantitative analysis of ^18^F-FDG PET uptake across different dosage levels was performed after normalizing regional data with a standard uptake value ratio (SUVr) to the cerebellum (Oblak et al., 2021;Onos et al., 2022). Using principal component analysis (PCA), twelve out of the twenty-seven studied regions that consistently explained 80% of the variance in ^18^F-FDG uptake between and within cohorts were identified (see Figure 2). In this model system, dynamic range was determined to have a 1.5-fold difference in ^18^F-FDG uptake between WT and 5XFAD mice (Oblak et al., 2021), thus supporting its use for drug discovery studies. Significance was seen at the brain region level across treatments (Onos et al., 2022), but pairwise interregional interactions were not explained by per-region significance, so a network approach (Jeub et al., 2018;Veronese et al., 2019;Chumin et al., 2023) was applied to further investigate interregional metabolic alterations.

**Figure 2.**
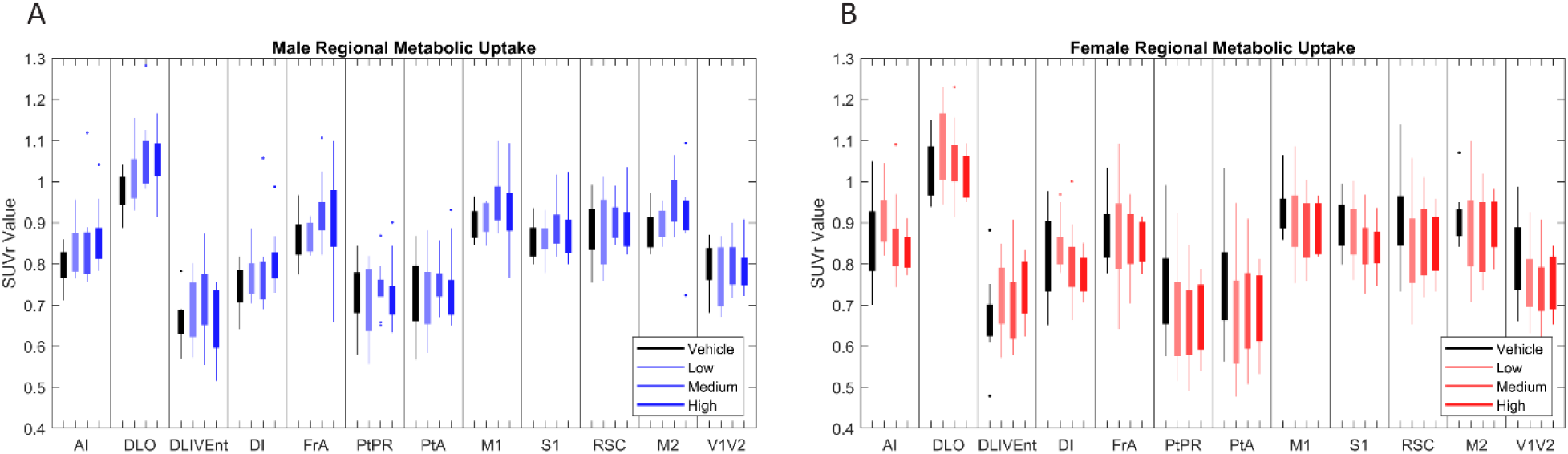
[18F]-FDG PET standardized uptake value ratio (SUVr) to cerebellum in brains regions in (A) male (N=39) and (B) female mice (N=38). Principle component analysis (PCA) found that the twelve brain regions above most consistently accounted for 80% of variance in cerebral metabolic uptake across cohorts. Full names of annotated brain region labels can be found in Supplementary Table 1.

**Figure 3.**
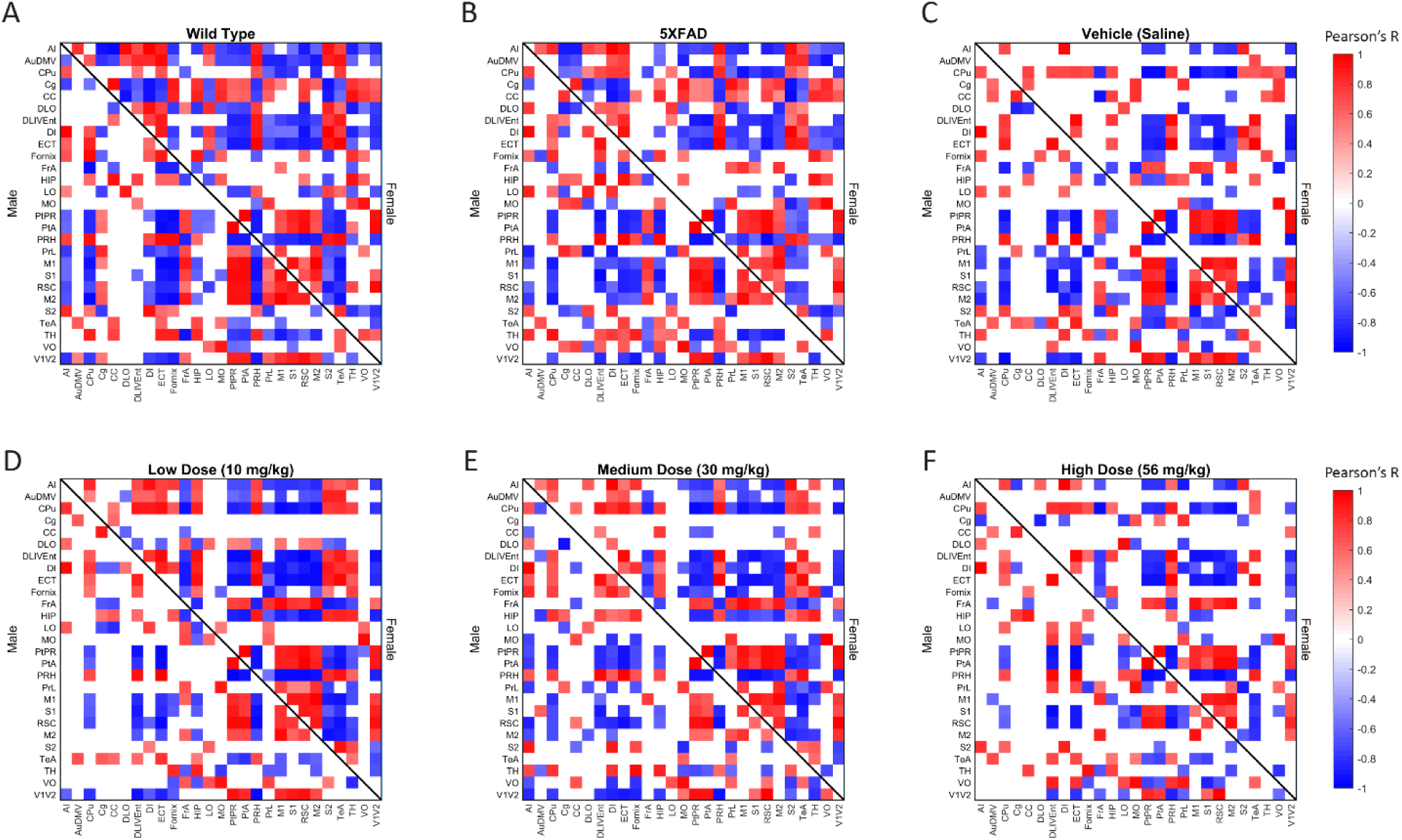
Thresholded metabolic covariance matrices sorted alphabetically by region. Edges were thresholded based on correlation magnitude, with surviving edges having a correlation significance value of p<0.05.

### Global Network Properties

**Figure 4.**
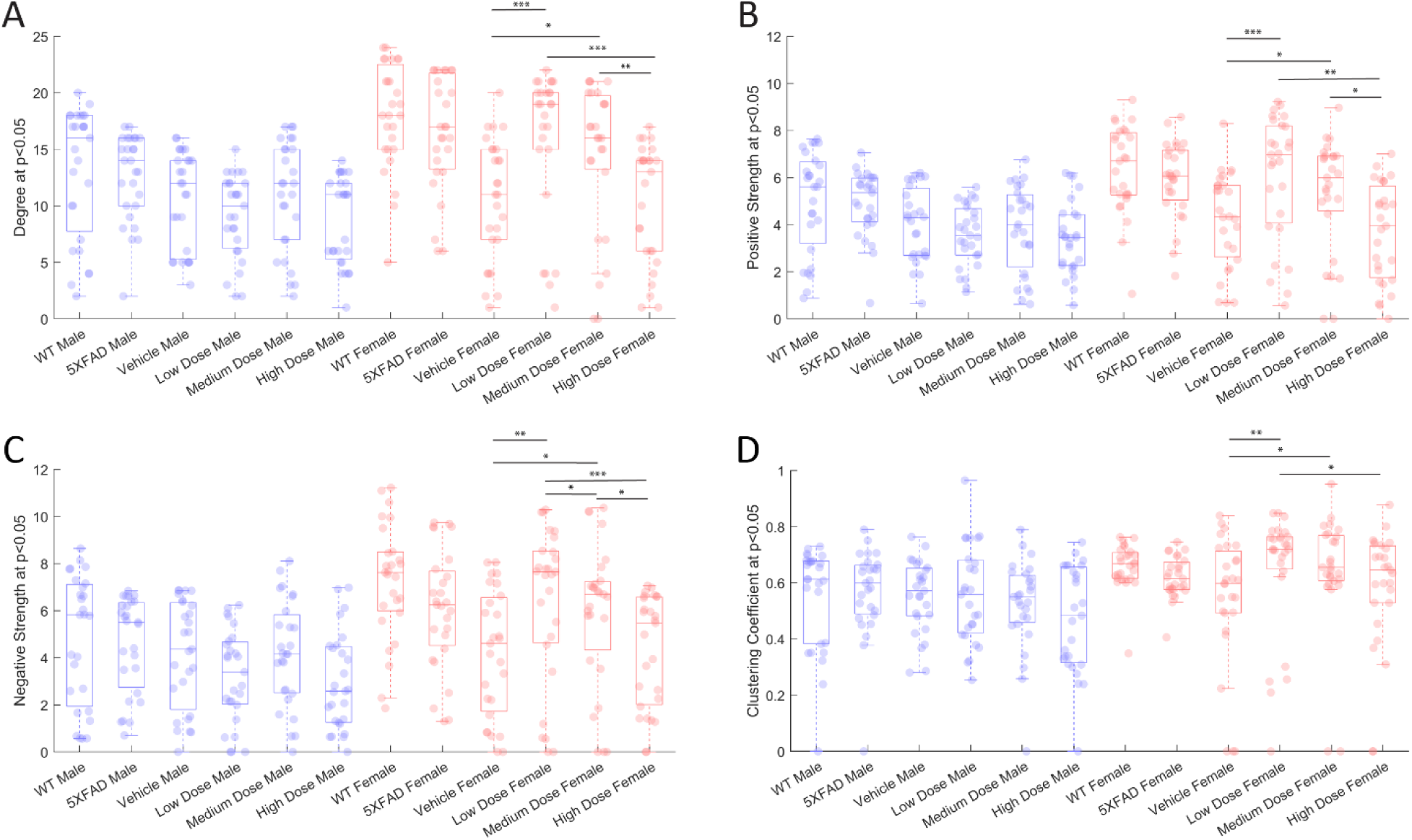
Properties of p<0.05 thresholded metabolic covariance for control and treatment groups. (A) Network degree, (B) positive strength, (C) negative strength, and (D) clustering coefficient. * denotes p < 0.05, ** denotes p<0.01, *** denotes p<0.005, 2-sample Kolmogorov-Smirnov test significance.

To measure metabolic changes at a network level, we utilized covariance and connectomic analyses (Jeub et al., 2018;Veronese et al., 2019;Chumin et al., 2023). First, we measured the degree of functional metabolic connectivity on a single-region basis, and extrapolated nodal degree values to a global distribution. For female mice, the network degree of vehicle dose and low dose (*p*<0.001), vehicle dose and medium dose (*p*<0.05), medium dose and high dose (*p*<0.01), and low dose and high dose (*p*<0.001) were significantly different. Interestingly, the vehicle dose and high dose had almost the same network distribution for degree (*p*=0.994), consistent with the findings of previous literature (Palop et al., 2007). By contrast, male mice did not follow the same degree changes observed in females, with no two distributions significantly different from one another between treatments (Figure 4A). Qualitatively, we see different connectivity patterns between sexes as a function of treatment. Males displayed oscillatory behavior in degree distributions, with the vehicle group more connected than low and high doses, but less connected than medium dose. By contrast, female mice showed a clear dose-dependent change in degree of connections, with the networks of low and medium dosed mice significantly more connected than those of the vehicle and high dosed mice. To look further into the weight and sign of functional connections between regions, we analyzed the positive and negative weighted node degree of every region in the brain, known and positive and negative strength, respectively. In females, the distributions of positive node strengths across the brain after a *p*<0.05 threshold was applied significantly differed between vehicle and low dose (*p*<0.001), vehicle and medium dose (*p*<0.05), medium and high dose (*p*<0.01), and low and high dose (*p*<0.01). Negative strengths in females resembled positive strengths in their distributions across treatment, both qualitatively in the same inverted paraboloid shape (see Figure 4B,C), and quantitatively with significant differences between the same groups as positive strengths, with the addition of significance between low and medium dose (*p*<0.05) not seen in positive strengths. Males did not show any significant differences between distributions of either positive or negative strengths after a p<0.05 threshold was applied to the covariance matrix. To measure the functional interconnectivity of brain subnetworks, clustering coefficients (propensity of connected triangles in a network) were calculated and compared in the same manner as the prior network characteristics. The clustering coefficients of brain networks followed a similar pattern across treatment for each sex. Significance was seen in the distribution of clustering coefficient in female networks between vehicle dose and low dose (*p*<0.01), vehicle dose and medium dose (*p*<0.05) and low dose and high dose (*p*<0.05), while males did not show significant differences that survived the *p*<0.05 threshold.

### Effect of Treatment on Covariance Distributions

Distributions of Pearson’s correlation between regions denoted by edge weight differed across sex and treatment, as measured by a two-sample KS non-parametric distribution comparison (see Figures 5-7). Males exhibited little handling stress, with qualitatively similar partitioned adjacency and edgewise distribution between 5XFAD disease control and saline-dosed vehicle cohorts (Figure 6B). Males treated chronically with a low dose (10 mg/kg) of LEV yielded a more normally distributed set of edgewise covariance trending towards significance compared to the vehicle (saline) treatment group of the same sex (KS = 0.01, *p* = 0.057). By contrast, males treated with the high dose (56 mg/kg) of LEV yielded a covariance distribution significantly different from the vehicle treated group (KS = 0.12, *p* < 0.05) (Figure 5E). Unlike males, females exhibited relationships in ^18^F-FDG covariance between treatments in the medium (30 mg/kg) and high (56 mg/kg) doses groups which trended towards significance (Figure 7D,E). Importantly, there was a significant sex effect with treatment distributions, where male and female cohorts were significantly different between low (KS = 0.15, *p* < 0.005) and medium (KS = 0.1, *p* < 0.05) dose, with males qualitatively more normally distributed than females (Figure 5).

**Figure 5.**
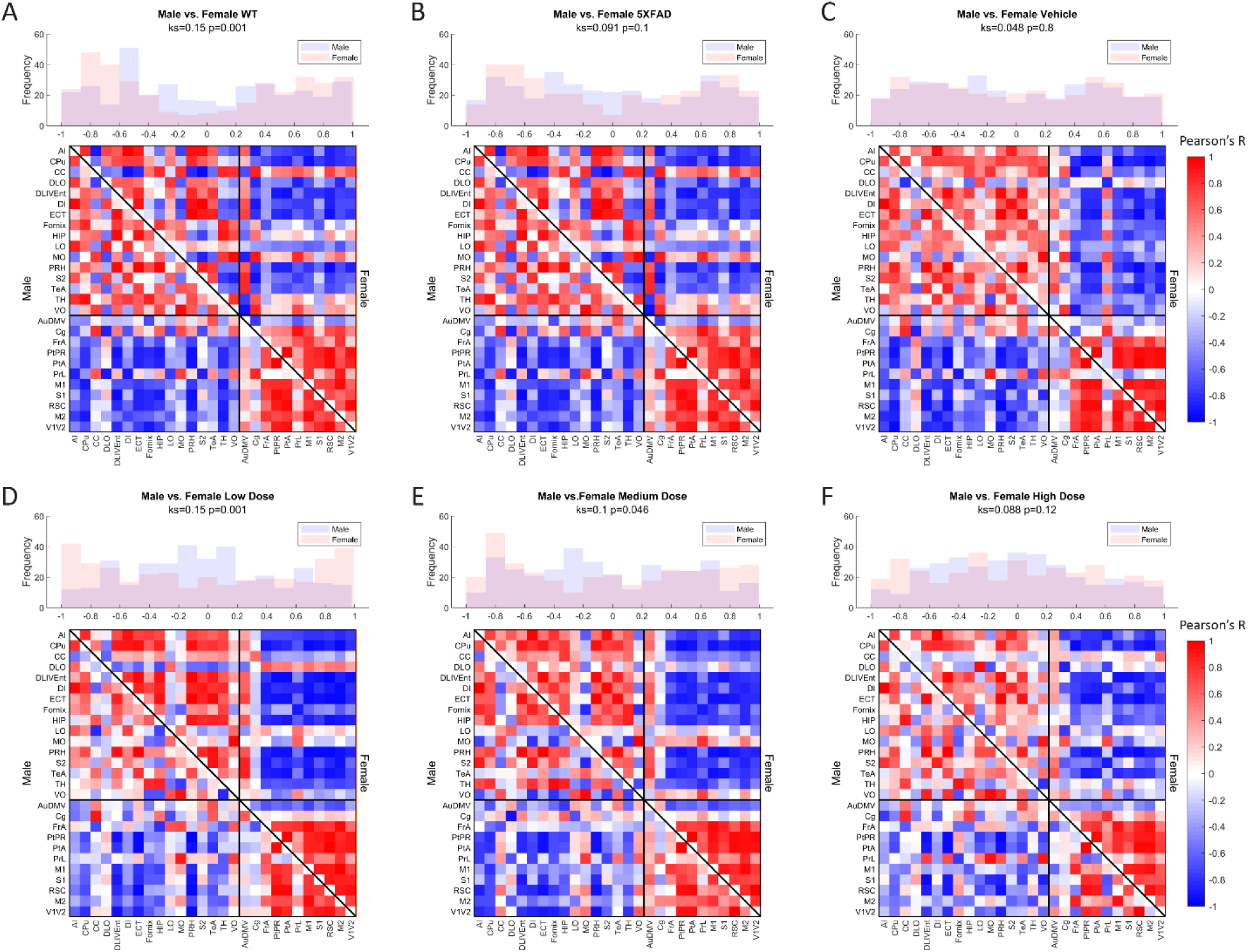
Metabolic ([18F]-FDG) covariance matrices for (A) wild-type (WT) (N=22, n=11 male, n=11 female), (B) 5XFAD (N=22, n=10 male, n=12 female), (C) saline vehicle dose (N=17, n=9 males, n=8 females), (D) low dose (N=21, n=10 males, n=11 females), (E) medium dose (N=20, n=10 males, n=10 females), and (F) high dose (N=19, n=10 males, n=9 females) treatment groups with male WT community partitions applied on upper (female) and lower (male) triangles. Histograms of interregional correlation values indicate correlation distribution across all brain regions for each respective cohort. Covariance was computed as Pearson’s correlation of z-scored regional SUVR values across animals within each group. Distributions between sexes of a given cohort are compared with a 2-sample Kolmogorov-Smirnov (KS) nonparametric test, with KS-value and p-value printed above the histogram.

**Figure 6.**
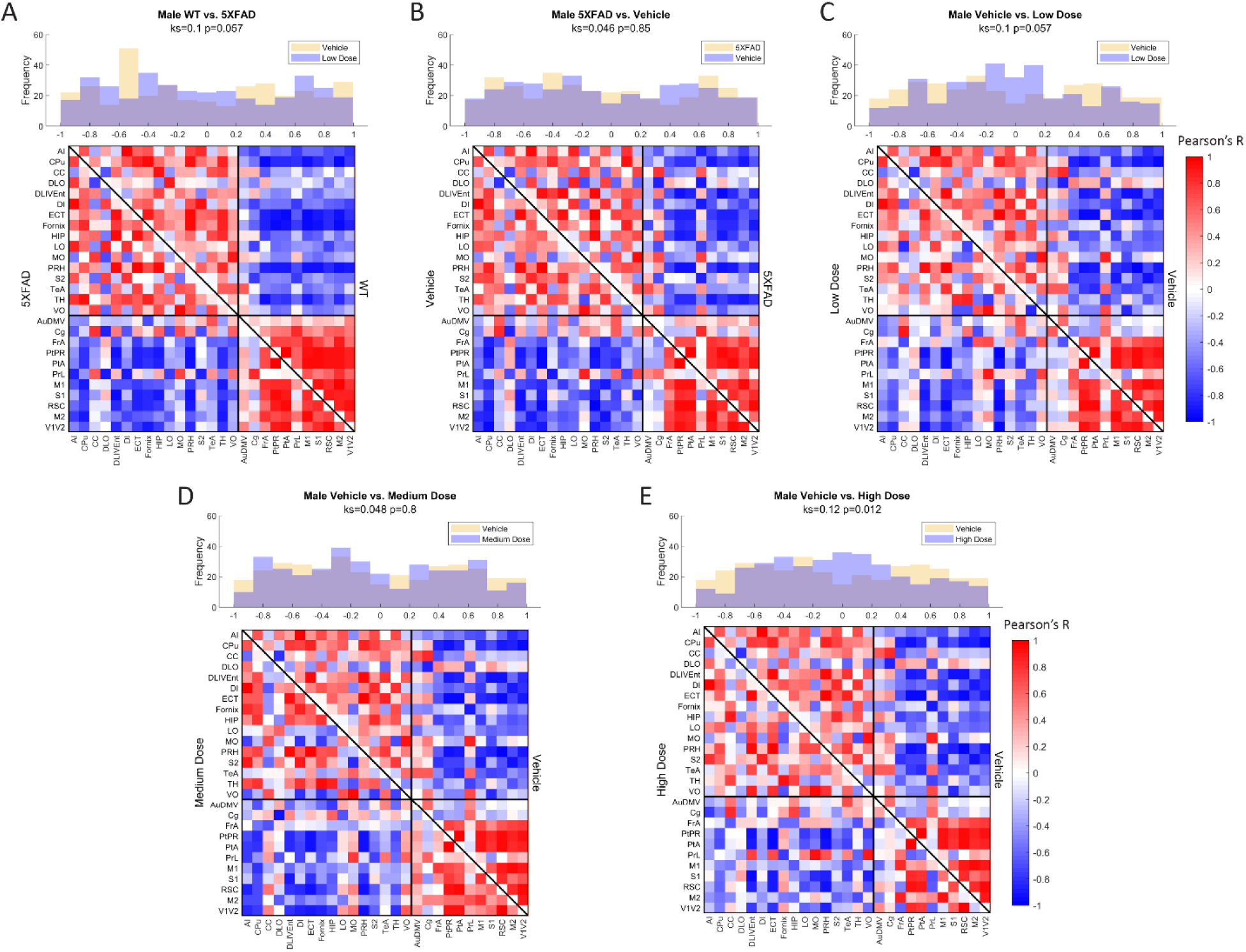
Male metabolic covariance matrices across treatment with WT community partitions applied. Changes in functional network structure can be observed via (A) changes in network and covariance distribution due to disease (N=21, n=11 WT, n=10 5XFAD), (B) handling stress (N=19, n=10 5XFAD, n=9 vehicle), (C) low (N=19, n=9 vehicle, n=10 low dose), (D) medium (N=19, n=9 vehicle, n=10 medium dose), and (E) high (N=19, n=9 vehicle, n=10 high dose) dose difference.

**Figure 7.**
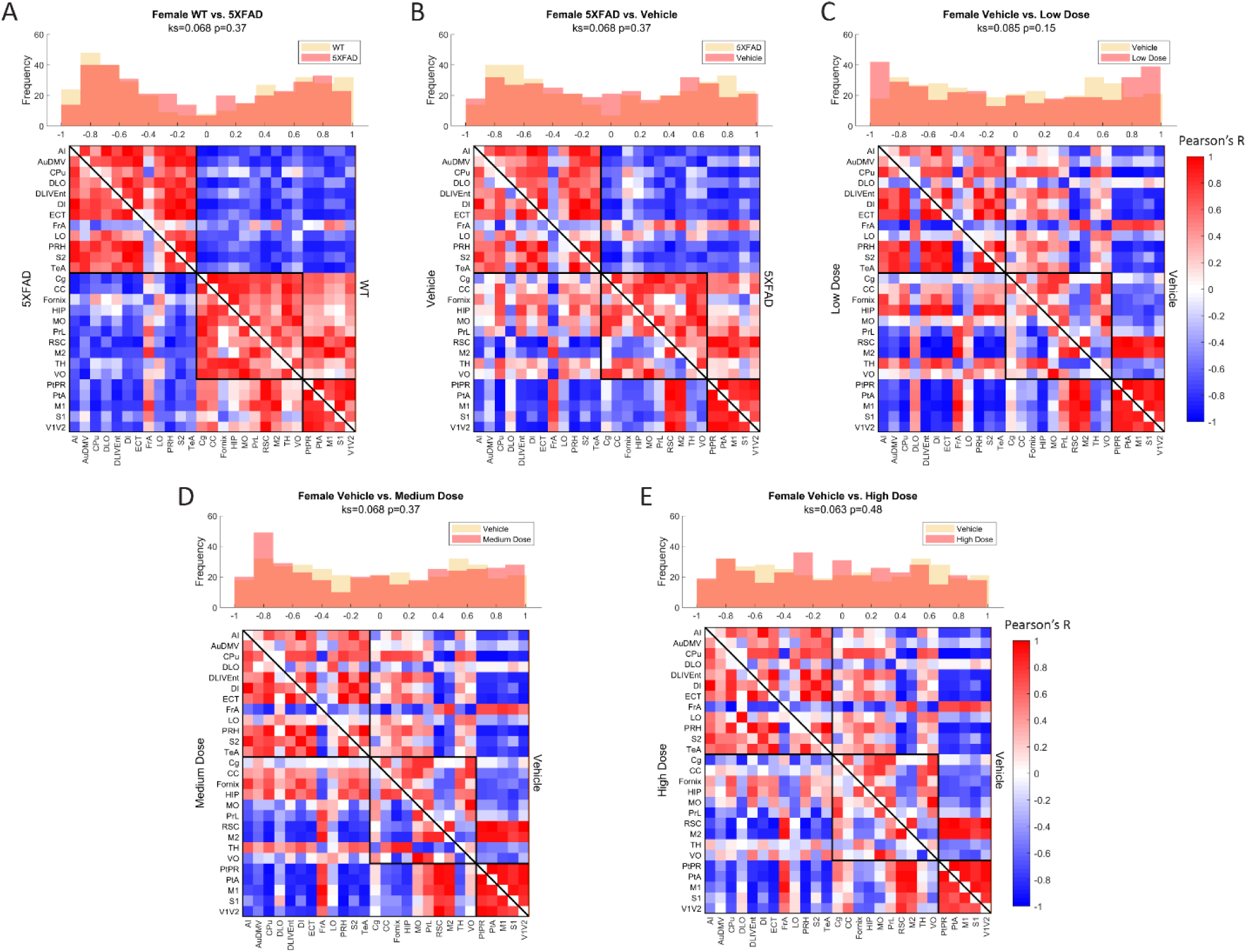
Female metabolic covariance matrices across treatment with WT community partitions applied. Changes in functional network structure can be observed via (A) changes in network and covariance distribution due to disease (N=23, n=11 WT, n=12 5XFAD), (B) handling stress (N=20, n=12 5XFAD, n=8 vehicle), (C) low (N=19, n=8 vehicle, n=11 low dose), (D) medium (N=18, n=8 vehicle, n=10 medium dose), and (E) high (N=17, n=8 vehicle, n=9 high dose) dose difference.

### Community Structure and Component Function with Sex

Metabolic covariance networks were assessed via the MRCC community detection algorithm with 10,000 permutations to establish a rigorous partition for covariance networks focused on positive covariance between regions (Veronese et al., 2019;Chumin et al., 2023) where partitions, or communities, differed between sexes and treatments. Female modularity ranged from two to five communities within a network, with the number of partitions generally increasing linearly as a function of dose. Male modularity consistently had more partitions relative to females, with the lowest number of modules present in the low dose group and an average of 5.66 modules across other treatments. We interpreted the number of communities as indicative of edgewise coherence among regions, akin to functional subnetworks. We observe that males generally exhibit a greater number of subnetworks relative to females. This agrees with our findings from the observations of node degree and strength distributions. Males form subnetworks of highly similar metabolic covariance, while the most similar partition structure for females tends towards large, bidirectional networks, consistent with higher degrees of connectivity across regions relative to males across treatments groups. To track network changes in a common reference space, we imposed the community structure of the healthy control WT male metabolic covariance network onto male and female network of the same treatment across all treatments (Figure 5). Male littermate controls were chosen as the reference network partition due to consensus findings of females displaying more aggressive AD disease phenotype at a given age, both in the 5XFAD model and in human AD pathology (Oblak et al., 2021). Using male littermate controls therefore enabled better tracking of deviations from a healthy metabolic network. To assess differences in metabolic uptake distributions in subnetworks between males and females at a given treatment, we performed ANOVA between male and female metabolic SUVr networks clustered by WT male MRCC partitions. Significance was corrected for multiple comparisons via Bonferroni correction. We found that at vehicle, low, and high dose, there were no significant differences in metabolic uptake in communities of the same regional groupings between sexes; however, at low dose, of the two communities detected in the male MRCC algorithm, one yielded significantly different metabolic uptake. The first community was composed of the following region members: AI, CPu, CC, DLO, DLIVEnt, DI, ECT, Fornix, HIP, LO, MO, PRH, S2, TeA, TH, and VO (See Supplemental Table 1 for full region names). At low dose, females exhibited significantly more uptake in this subnetwork than males (F = 4.9, *p*<0.05). The functional interpretation of this subnetwork, using equally weighted components, was determined to be primarily learning, with additional components of perception and sensory processing. Within sex, males displayed significantly different metabolic uptake between vehicle and medium dose in the subnetwork containing AuDMV, Cg, FrA, PtPR, PtA, PrL, M1, S1, RSC, M2, and V1V2 (F = 4.7, *p*<0.05). Functionally, the regions within this subnetwork encoded primarily for learning and sensory processing. Females did not display significantly different uptake in common communities between treatments.

### nanoString Gene Expression Profiling

Brain hemispheres were assessed via nanoString for gene expression differences using a panel based on human AD gene expression changes (Allen et al., 2016;Mostafavi et al., 2018;Wang et al., 2018;Preuss et al., 2020;Wan et al., 2020). Linear regression analysis revealed 15 genes that were significant at *p*<0.05 for genotype, sex, and/or treatment. As shown in Figure 8A, the change in log2 intensities from WT vehicle mice demonstrated undirected significant dose-specific effects for genes which encode for AD, epileptiform activity, cerebral metabolism, and/or cerebral homeostatic mechanisms. Further analyses were performed on genes from medium and high dose treated animals that were significantly anti-correlated with neuronal-related human modules in the inferior frontal gyrus (IFG) region (Wan et al., 2020). This gene module identified 129 genes, and Gene ontology (GO) enrichment analysis identified a number of biological processes, which, consistent with the mechanism of action of LEV (Dubovsky et al., 2015), included synaptic vesicle cycling, vesicle exocytosis, vesicle recycling, vesicle priming, regulated exocytosis, neurotransmitter transport, neurotransmitter secretion, and protein localization to cellular junctions (Wan et al., 2020;Onos et al., 2022) (see Figure 8B). Further pathway analyses performed on these 129 genes with Ingenuity Pathway Analysis (IPA) (Kramer et al., 2014) revealed significant overlap with canonical pathways ‘SNARE signaling pathway’, ‘Netrin Signaling’, ‘Synaptogenesis signaling pathway’, ‘mitochondrial dysfunction’ and ‘Glutaminergic Receptor Signaling pathway (enhanced)’. Top regulator effect networks were also identified, such as’ ‘Exocytosis and Learning’ comprised of *BDNF*, *FMR1*, *NFE2L2* and *PHF21A*, and ‘Seizures’ comprised of *DNMT3A*, *PIAS1* and *REST*. Lastly, ‘memantine’ was determined to be both a significant upstream regulator and driver of the causal network. Memantine (Lipton, 2006), also known as Namenda, was one of the first approved drugs to treat AD and has a similar target to LEV: it works as an NMDA receptor antagonist aimed at quelling abnormal activity in the brain. Overall, identification of specific processes and pathways relevant to drug treatment help to narrow mechanisms and refine preclinical translation from mouse to human studies.

**Figure 8.**
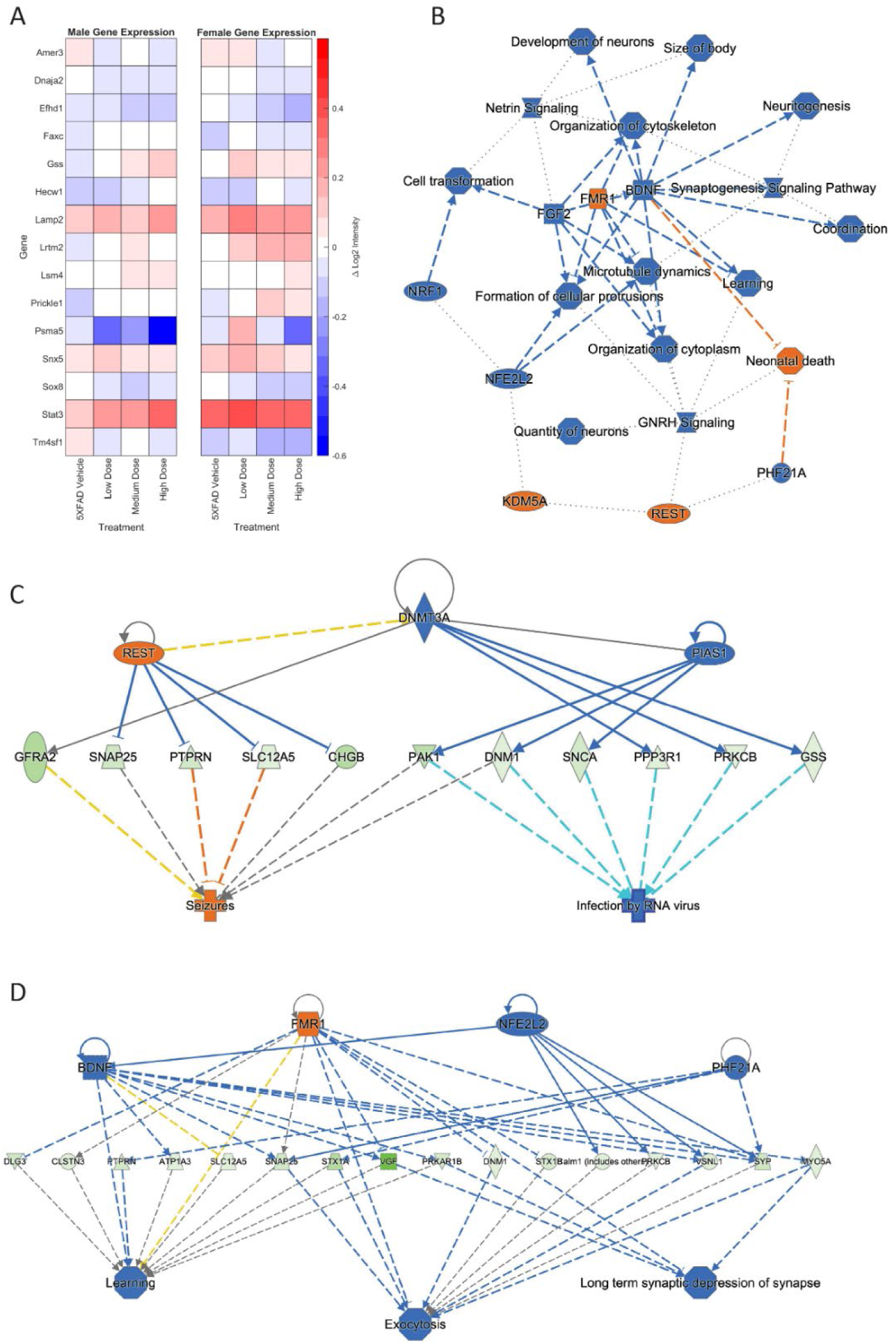
Hemisagittal mouse brain sections were homogenized and analyzed through nanoString. (A) Log2 intensity gene expression in male (n=39) and female (n=38) 5XFAD treatment groups from male and female WT vehicle groups, respectively. Gene expression at treatment level was subtracted from expression at vehicle, and results plotted as a heat map. Genes plotted are linked to AD, epileptiform activity, cerebral metabolism, and/or cerebral homeostatic mechanisms. (B) Primary GO pathway analysis found the processes connected to similarly expressed genes between medium and high dosed female mice. Further pathway analysis focused found connections to (C) seizure, (D) learning, exocytosis, and long-term depression functions. See Supplementary figure 1 for prediction legend.

### Exposure-Response Modeling

Despite the plasma concentration AUC relationship and the dose dependent changes observed with metabolic connectomics, no apparent relationships could be established with connectomics data when PK/PD modelled using a multi-linear regression. By contrast, expression changes in Amer3, Lamp2, Lrtm2, Psma5, and Stat3 all showed significant associations (*p*<0.05) with LEV concentration and/or animal sex (Table 1, Figure 9).

**Figure 9.**
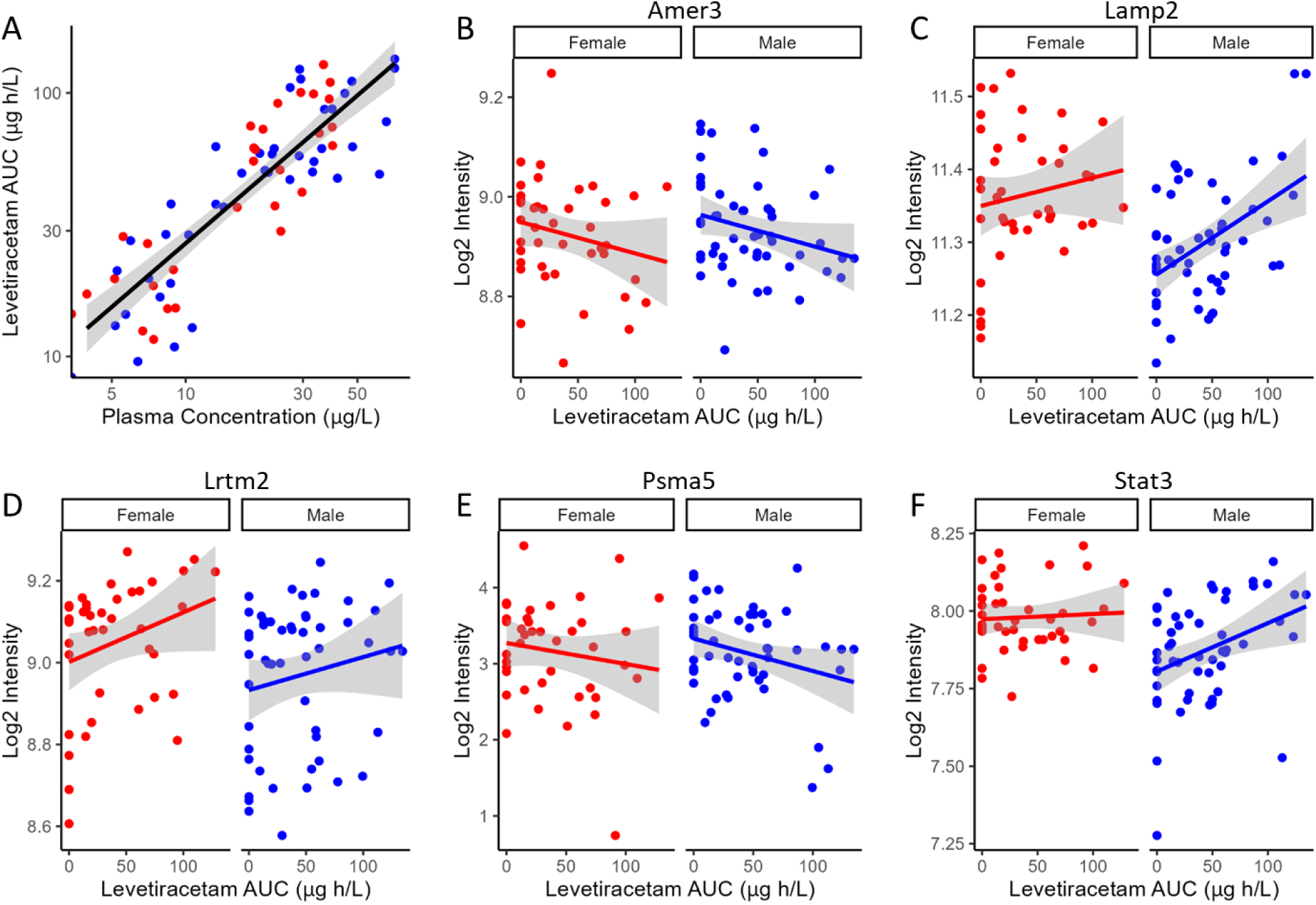
PK/PD plots of Log2 Intensity as a function of levetiracetam exposure for (A) Amer3, (B) Lamp2, (C) Lrtm2, (D) Psma5, and (E) Stat3 expression are associated with predicted levetiracetam exposure (AUC) using a linear PK/PD model. Solid red (female) and blue (male) lines indicate linear regression, while shaded region represent the 95% confidence interval.

**Table 1.** Association between gene expression and levetiracetam exposure area under the curve (AUC) and sex. Modelling was calculated in R using lm(Y∼AUC+SEX).

| | AUC $\beta \pm$ SE (p-value) | Sex $\beta \pm$ SE (p-value) | R<sup>2</sup> |
| --- | --- | --- | --- |
| Amer3 | -0.006 ± 0.0003 (0.024) | 0.014 ± 0.021 (0.5) | 0.04 |
| Lamp2 | 0.00075 ± 0.0002 (0.0013) | -0.069 ± 0.017 (<0.001) | 0.21 |
| Lrtm2 | 0.00098 ± 0.00046 (0.04) | -0.085 ± 0.035 (0.02) | 0.078 |
| Psma5 | -0.0036 ± 0.0017 (0.04) | -0.0016 ± 0.13 (0.99) | 0.025 |
| Stat3 | 0.001 ± 0.0004 (0.16) | -0.11±0.03 (<0.001) | 0.16 |

## DISCUSSION

In the current study, we performed retrospective network analysis of LEV data to gain novel insight about how chronic dosing affects AD progression as it pertains to cerebral metabolic uptake and gene expression. Our results show that 6-month-old 5XFAD mice display significant changes in metabolic uptake as seen by network analyses correlated within and between regions in both sexes and across treatment. Per previous studies, we conducted general linear statistical modeling of regional ^18^F-FDG uptake data as a measure of regional brain function. Quantitative analysis of these data revealed a sex-by-dose relationship (Onos et al., 2022); however, due to limitations of these approaches, post-hoc analysis only permitted pair-wise analysis of regional data, thus negating the inter-regional network changes that support higher brain function (Chumin et al., 2023). To overcome this, we applied network covariance analysis to ^18^F-FDG data, which has been shown to distinguish stage of disease for both preclinical (Chumin et al., 2023) and clinical AD (Veronese et al., 2019). Using this approach, we quantified pairwise network covariance for all regions, then computed edgewise network features by calculating distributions of degree, positive and negative strength, and clustering coefficient in metabolic networks from thresholded networks. We extrapolated per-region node characteristics to global distributions and compared changes in distributions between treatment within sex. At 6 months, female mice displayed significantly more interregional metabolic uptake as seen by the degree and density of both diseased mice and their littermate controls (Figure 4). Overall, network characteristics indicated that the magnitude of interregional metabolic correlation decreased as chronic dose concentration increased in males, as seen in global degree, and strength outputs. In females, disease and handling stress decreased interregional functional connectivity, some of which was recovered in a dose-dependent manner. Females dosed chronically at 10 mg/kg showed the greatest metabolic network change across analyses, consistent with the mode of action of LEV. More generally, network alteration was inversely proportional to dose concentration in females (Figure 4).

Differences in uptake in control cohorts were not localized to specific communities between sexes in 5XFAD mice when the male community partition was applied to both groups, but differences were localized to module 1 of the male partition structure in WT mice (*p*<0.05, Bonferroni corrected). While vehicle, medium, and high doses did not exhibit significantly different uptake between like communities, low dose male and female mice exhibited significant differences in metabolic uptake in one of two modules. Based on the region members of this module, these changes are expected to impact learning, memory, and sensory integration, and are consistent with LEV mode of action.

5XFAD is known to be an aggressive amyloidogenic early-onset model of Alzheimer’s disease. Like AD, 5XFAD mice exhibit sexual dimorphism in pathology, with females displaying a more rapid onset disease phenotype (Oblak et al., 2021). Previous studies have presented conflicting findings on cerebral metabolic uptake in the 5XFAD model around 6 months (Oblak et al., 2021). In the current study, we treat littermate controls as a reference population and explore the changes of metabolic networks with respect to littermate controls. We hypothesized that within sex, we would see a dose-dependent response evident in network characteristics and community structure. We also hypothesized that this dose-dependency would vary between sexes, with females showing a greater restorative response in part due to their faster progression towards disease state.

Comparing littermate controls to diseased (5XFAD) mice, there were surprisingly few differences in male and female adjacency matrices when community partition structure of their WT littermates was applied. By contrast, female 5XFAD mice exhibited a greater handling stress response than males, and this stress response was localized to interregional correlation shifts in the retrosplinial cortex (RSC) and secondary motor cortex (M2). This is consistent with amyloid deposition previously reported for these structures (Oblak et al., 2021). This handling effect was observed in all female LEV dosed cohorts (see Figure 7B-E). At low dose, neither females nor males had network alterations significantly different from vehicle, though males trended towards significance. At medium dose, females showed qualitative differences in network structure compared to vehicle in the interregional correlations of the frontal association cortex as well as the lateral orbitofrontal cortex. Males showed similar qualitative differences in lateral orbitofrontal correlations, but neither sex differed from vehicle at the medium dose. At high dose, males significantly differed from vehicle in edgewise correlations with many weak correlations relative to vehicle (*p*<0.05) (Figure 6E). Females, on the other hand, did not display significant edgewise distributions, and instead converged back towards vehicle distribution relative to medium and low dose. This finding is consistent with dose-dependent findings of previous LEV studies (Onos et al., 2022); at high chronic doses of LEV, females both in mice as well as in the clinic develop a functional resistance to the drug, the exact cause of which is yet to be uncovered (Löscher and Schmidt, 2006;Surges et al., 2008;Sanchez et al., 2012;Onos et al., 2022).WT covariance distributions were significantly different between males and females (*p*<0.005). Females display a greater number of strong correlations, both positive and negative, between regions. This is consistent with previous literature showing that WT females show marginally greater glycolytic metabolism in the brain after ∼6 months of age (Oblak et al., 2021). 5XFAD mice displayed no significant difference in their distributions, however, females did show a bimodal distribution with the highest frequency of covariance values occurring at the limits. In low and medium doses, female edgewise distributions were again bimodal and significantly different from males (*p*<0.005 and *p*<0.05, respectively). These differences further reinforce the sexual dimorphism of 5XFAD mice which has been previously reported (Oblak et al., 2021;Onos et al., 2022;Chumin et al., 2023;Jullienne et al., 2023), and are thought to be linked to the transgene expression being linked to the estrogen sensitive Thy1 promoter (Moechars et al., 1996).

Network analyses of thresholded metabolic correlation networks, like edgewise distributions, revealed network characteristics that differed on a sex by treatment basis. WT mice follow a pattern with females’ retaining greater numbers of nodes than their male counterpart, as seen in network degree and clustering coefficient distributions (Figure 3A,D). Vehicle cohorts showed a sex-specific decrease in density relative to WT and likely represents handling stress. This handling stress is consistent with prior findings which show that female mice exhibit higher stress hormone response than their male counterparts when introduced to handling stressors (Sensini et al., 2020). At low and medium dose, we see network degree approach WT levels only to decline at a high dose of LEV, which we interpreted as due to functional tolerance. Further analysis of positive and negative network strengths support the sex and dose dependency in whole brain networks. Females contain more positive and negative strengths than males at every treatment, and the low and medium LEV doses show a significant increase in positive and negative strengths (Figure 4B,C). This supports the notion that in AD pathology, signal disruption is seen both in suppression of some circuits and aberrant activity of others, resulting in more extreme interregional relationships (Palop et al., 2007). Previous studies (Palop, 2009;Sanchez et al., 2012;Onos et al., 2022) have reported that LEV reduces sub-seizure epileptiform activity in mouse models of AD (Chumin et al., 2023) and we believe that by reducing aberrant neuronal activity, LEV perturbs hyperactive neural circuitry back towards a functional equilibrium with decreased signal suppression distal to Aβ plaques and a regulation of aberrant activity near plaques. However, specificity between equilibrium metabolic uptake in neurons, glia, and other energy-demanding cerebral machinery and nonequilibrium inflammation-driven metabolic uptake in microglia and hyperactive dysregulated circuits cannot be distinguished in ^18^F-FDG PET imaging (Xiang et al., 2021). Because we were unable to empirically differentiate between cerebral metabolic contributors due to the retrospective nature of the current study, we conducted targeted transcriptomic analysis using nanoString to track gene expression and identified fifteen genes to be dose and sex dependent (See Figure 8A). Further analyses examining GO biological processes and pathway overlap in genes shared between medium and high-dosed animals were primarily representative of synaptic function and organization as it pertains to neural activity. REST gene activation, whose pathway causes suppression of genes related to epileptiform activity and seizures was identified as a pathway, as was the FMR1 pathway, implicated in the activation of genes associated with synaptic depression and learning (see Figure 8C and D, respectively), which is consistent with LEV’s mode of action. Together, these pathways contribute to a larger transcriptional network at play in mice treated with LEV composed primarily of neuronal maintenance and restoration (Figure 8B). This was not overly surprising, as the genes analyzed were selected as they were correlated with human AD genes in the Neuronal Consensus cluster. However, connecting gene pathways to metabolic uptake helped to link cerebral metabolic allocation corresponding to LEV treatment in a way that is currently unattainable by PET alone.

As with all studies, there are limitations that must be considered when drawing conclusions. First, the 5XFAD model was developed as a rapid-onset, highly aggressive representation of early-onset Alzheimer’s disease. 5XFAD displays early amyloid beta plaque formation relative to other mouse models and epileptiform activity localized to the dentate gyrus and hippocampus, spatially analogous to human data (Abe et al., 2020). However, 5XFAD mice have been shown not to express behavioral disease phenotype relative to controls analogous to MCI and AD patients in the clinic (Oblak et al., 2021). With this in mind, the current study revealed novel interactions that can provide insight into cerebral metabolic modulation by LEV in AD. LEV clearly acts in a dose and sex-dependent manner, with network alteration relying on the degree of disease phenotype present in the subject. To more completely understand the network effects of LEV, future studies might use less severe, more physiologically analogous mouse models of AD, and bear in mind sexual dimorphism to test LEV efficacy in both sexes at equivalent disease phenotypes. MODEL-AD is currently longitudinally testing novel models of AD, which may fulfill these requirements (Oblak et al., 2020)] and studying the effect of LEV on disease progression in these models may be more translationally viable. An additional limitation discussed in the original study is the choice of LEV dose chronically administered to these mice (Onos et al., 2022). Future studies looking at LEV’s metabolic modulatory effects may choose more intermediate doses than the ones in this study gain a more in-depth understanding of edgewise changes in community structures as well as between regions. Analytical tools to connect regional components of functional metabolic correlation communities to their emergent functional properties such as sensory, motor, or integration hubs are currently in development, and will shed light on the nature of metabolic restructuring in terms of edgewise correlation shifts.

Previous studies show that LEV likely acts near circuits or networks plagued by beta amyloid deposition and that behavioral alterations can be tracked in a dose-dependent manner (Busche et al., 2008;Sanchez et al., 2012). In the original PTC study, mice underwent ^18^F-AV45 PET imaging according to the same protocols of the ^18^F-FDG imaging as well as behavioral evaluation (Onos et al., 2022). A lack of significant change in behavioral and amyloid load differences led us to exclude these data in our retrospective analysis. However, future studies utilizing PET imaging of amyloid deposition in mouse models to map the co-localization of this structural network with the functional networks given by metabolic PET imaging may shed more insight into metabolic perturbation of AD pathology with respect to amyloid imaging. Moreover, combining these with transcriptomics and PK/PD modeling could lead to a more precise indication of drug efficacy. This, in turn, would have significant translational implications for precision medicine and treatment of the synchronized hyperactive neural activity seen in patients with AD.

In the previous study, gene expression analysis revealed that in an Accelerating Medicines Partnership Program for Alzheimer’s Disease (AMP AD) consensus clusters of genes, females exhibit anticorrelation in gene expression with respect to analogous human expression, suggesting that at high chronic dosing of LEV, there may be effects seen that cannot be explained by our functional tolerance interpretation (Preuss et al., 2020;Onos et al., 2022). Future work may utilize a multimodal approach involving higher throughput and/or spatial transcriptomics coupled with metabolic uptake quantification to better resolve ambiguities in the physiological effects of LEV dose-wise.

In conclusion, our research revealed significant dose-dependent changes in region-wise glycolytic correlation and gene expression changes in response to LEV not possible with traditional general linear statistical modeling. Further, it revealed a sex and dose dependency that provides a means to track network changes using translationally relevant ^18^F-FDG PET imaging.

## Data Availability

All protocols, raw, and summary data are available at https://adknowledgeportal.org, where the Synapse ID is syn2580853, and DOI is 10.7303/syn2580853.

## Acknowledgements

The authors would like to acknowledge the JAX Center for Biometric Analysis staff for support of the chronic dosing studies. In addition, we would like to acknowledge the JAX Genome Technologies core for assistance with nanoString RNA extractions and sample processing. The current study data, in whole or in part, were obtained from the AD Knowledge Portal (https://adknowledgeportal.org), and the brain covariance analysis software was obtained from the publicly available CovNet package (https://github.com/echumin/CovNet).

## References

Abe, Y., Ikegawa, N., Yoshida, K., Muramatsu, K., Hattori, S., Kawai, K., Murakami, M., Tanaka, T., Goda, W., Goto, M., Yamamoto, T., Hashimoto, T., Yamada, K., Shibata, T., Misawa, H., Mimura, M., Tanaka, K.F., Miyakawa, T., Iwatsubo, T., Hata, J.-I., Niikura, T., and Yasui, M. (2020). Behavioral and electrophysiological evidence for a neuroprotective role of aquaporin-4 in the 5xFAD transgenic mice model. Acta Neuropathologica Communications 8.

Allen, M., Carrasquillo, M.M., Funk, C., Heavner, B.D., Zou, F., Younkin, C.S., Burgess, J.D., Chai, H.S., Crook, J., Eddy, J.A., Li, H., Logsdon, B., Peters, M.A., Dang, K.K., Wang, X., Serie, D., Wang, C., Nguyen, T., Lincoln, S., Malphrus, K., Bisceglio, G., Li, M., Golde, T.E., Mangravite, L.M., Asmann, Y., Price, N.D., Petersen, R.C., Graff-Radford, N.R., Dickson, D.W., Younkin, S.G., and Ertekin-Taner, N. (2016). Human whole genome genotype and transcriptome data for Alzheimer’s and other neurodegenerative diseases. Sci Data 3, 160089.

Angulo, S.L., Henzi, T., Neymotin, S.A., Suarez, M.D., Lytton, W.W., Schwaller, B., and Moreno, H. (2020). Amyloid pathology-produced unexpected modifications of calcium homeostasis in hippocampal subicular dendrites. Alzheimer’s & Dementia 16, 251–261.

Bakker, A., Albert, M.S., Krauss, G., Speck, C.L., and Gallagher, M. (2015). Response of the medial temporal lobe network in amnestic mild cognitive impairment to therapeutic intervention assessed by fMRI and memory task performance. Neuroimage-Clinical 7, 688–698.

Bakker, A., Gregory, Marilyn, Caroline, Lauren, Craig, Michael, Susan, Amy, and Gallagher, M. (2012). Reduction of Hippocampal Hyperactivity Improves Cognition in Amnestic Mild Cognitive Impairment. Neuron 74, 467–474.

Busche, M.A., Eichhoff, G., Adelsberger, H., Abramowski, D., Wiederhold, K.H., Haass, C., Staufenbiel, M., Konnerth, A., and Garaschuk, O. (2008). Clusters of hyperactive neurons near amyloid plaques in a mouse model of Alzheimer’s disease. Science 321, 1686–1689.

Chumin, E., Burton, C., Silvola, R., Miner, E., Persohn, S., Veronese, M., and Territo, P. (2023). Brain Metabolic Network Covariance and Aging in a Mouse Model of Alzheimer’s Disease. bioRxiv, 2023.2006.2021.545918.

Dubovsky, S.L., Daurignac, E., Leonard, K.E., and Serotte, J.C. (2015). Levetiracetam, Calcium Antagonism, and Bipolar Disorder. J Clin Psychopharmacol 35, 422–427.

García-Pérez, E., Mahfooz, K., Covita, J., Zandueta, A., and Wesseling, J.F. (2015). Levetiracetam accelerates the onset of supply rate depression in synaptic vesicle trafficking. Epilepsia 56, 535–545.

Jeub, L.G.S., Sporns, O., and Fortunato, S. (2018). Multiresolution Consensus Clustering in Networks. Scientific Reports 8.

Jullienne, A., Szu, J.I., Quan, R., Trinh, M.V., Norouzi, T., Noarbe, B.P., Bedwell, A.A., Eldridge, K., Persohn, S.C., Territo, P.R., and Obenaus, A. (2023). Cortical cerebrovascular and metabolic perturbations in the 5xFAD mouse model of Alzheimer’s disease. Front Aging Neurosci 15, 1220036.

Klee, J.L., Kiliaan, A.J., Lipponen, A., and Battaglia, F.P. (2020). Reduced firing rates of pyramidal cells in the frontal cortex of APP/PS1 can be restored by acute treatment with levetiracetam. Neurobiology of Aging 96, 79–86.

Kramer, A., Green, J., Pollard, J., Jr., and Tugendreich, S. (2014). Causal analysis approaches in Ingenuity Pathway Analysis. Bioinformatics 30, 523–530.

Kumar, A., Maini, K., and Kadian, R. (2023). “Levetiracetam,” in StatPearls. (Treasure Island (FL)).

Lippa, C.F., Rosso, A., Hepler, M., Jenssen, S., Pillai, J., and Irwin, D. (2010). Levetiracetam: a practical option for seizure management in elderly patients with cognitive impairment. Am J Alzheimers Dis Other Demen 25, 149–154.

Lipton, S.A. (2006). Paradigm shift in neuroprotection by NMDA receptor blockade: memantine and beyond. Nat Rev Drug Discov 5, 160–170.

Löscher, W., and Schmidt, D. (2006). Experimental and Clinical Evidence for Loss of Effect (Tolerance) during Prolonged Treatment with Antiepileptic Drugs. Epilepsia 47, 1253–1284.

Lynch, B.A., Lambeng, N., Nocka, K., Kensel-Hammes, P., Bajjalieh, S.M., Matagne, A., and Fuks, B. (2004). The synaptic vesicle protein SV2A is the binding site for the antiepileptic drug levetiracetam. Proceedings of the National Academy of Sciences 101, 9861–9866.

Magalhães, J.C., Gongora, M., Vicente, R., Bittencourt, J., Tanaka, G., Velasques, B., Teixeira, S., Morato, G., Basile, L.F., Arias-Carrión, O., Pompeu, F.a.M.S., Cagy, M., and Ribeiro, P. (2015). The Influence of Levetiracetam in Cognitive Performance in Healthy Individuals: Neuropsychological, Behavioral and Electrophysiological Approach. Clinical Psychopharmacology and Neuroscience 13, 83–93.

Minkeviciene, R., Rheims, S., Dobszay, M.B., Zilberter, M., Hartikainen, J., Fülöp, L., Penke, B., Zilberter, Y., Harkany, T., Pitkänen, A., and Tanila, H. (2009). Amyloid β-Induced Neuronal Hyperexcitability Triggers Progressive Epilepsy. The Journal of Neuroscience 29, 3453–3462.

Moechars, D., Lorent, K., De Strooper, B., Dewachter, I., and Van Leuven, F. (1996). Expression in brain of amyloid precursor protein mutated in the alpha-secretase site causes disturbed behavior, neuronal degeneration and premature death in transgenic mice. EMBO J 15, 1265–1274.

Mostafavi, S., Gaiteri, C., Sullivan, S.E., White, C.C., Tasaki, S., Xu, J., Taga, M., Klein, H.U., Patrick, E., Komashko, V., Mccabe, C., Smith, R., Bradshaw, E.M., Root, D.E., Regev, A., Yu, L., Chibnik, L.B., Schneider, J.A., Young-Pearse, T.L., Bennett, D.A., and De Jager, P.L. (2018). A molecular network of the aging human brain provides insights into the pathology and cognitive decline of Alzheimer’s disease. Nat Neurosci 21, 811–819.

Musaeus, C.S., Shafi, M.M., Santarnecchi, E., Herman, S.T., and Press, D.Z. (2017). Levetiracetam Alters Oscillatory Connectivity in Alzheimer’s Disease. Journal of Alzheimers Disease 58, 1065–1076.

Oblak, A.L., Forner, S., Territo, P.R., Sasner, M., Carter, G.W., Howell, G.R., Sukoff-Rizzo, S.J., Logsdon, B.A., Mangravite, L.M., Mortazavi, A., Baglietto-Vargas, D., Green, K.N., Macgregor, G.R., Wood, M.A., Tenner, A.J., Laferla, F.M., Lamb, B.T., And The, M.-A., and Consortium (2020). Model organism development and evaluation for late-onset Alzheimer’s disease: MODEL-AD. Alzheimers Dement (N Y) 6, e12110.

Oblak, A.L., Lin, P.B., Kotredes, K.P., Pandey, R.S., Garceau, D., Williams, H.M., Uyar, A., O’rourke, R., O’rourke, S., Ingraham, C., Bednarczyk, D., Belanger, M., Cope, Z.A., Little, G.J., Williams, S.G., Ash, C., Bleckert, A., Ragan, T., Logsdon, B.A., Mangravite, L.M., Sukoff Rizzo, S.J., Territo, P.R., Carter, G.W., Howell, G.R., Sasner, M., and Lamb, B.T. (2021). Comprehensive Evaluation of the 5XFAD Mouse Model for Preclinical Testing Applications: A MODEL-AD Study. Front Aging Neurosci 13, 713726.

Onos, K.D., Quinney, S.K., Jones, D.R., Masters, A.R., Pandey, R., Keezer, K.J., Biesdorf, C., Metzger, I.F., Meyers, J.A., Peters, J., Persohn, S.C., Mccarthy, B.P., Bedwell, A.A., Figueiredo, L.L., Cope, Z.A., Sasner, M., Howell, G.R., Williams, H.M., Oblak, A.L., Lamb, B.T., Carter, G.W., Rizzo, S.J.S., and Territo, P.R. (2022). Pharmacokinetic, pharmacodynamic, and transcriptomic analysis of chronic levetiracetam treatment in 5XFAD mice: A MODEL-AD preclinical testing core study. Alzheimers Dement (N Y) 8, e12329.

Palop, J.J. (2009). Epilepsy and Cognitive Impairments in Alzheimer Disease. Archives of Neurology 66, 435.

Palop, J.J., Chin, J., Roberson, E.D., Wang, J., Thwin, M.T., Bien-Ly, N., Yoo, J., Ho, K.O., Yu, G.-Q., Kreitzer, A., Finkbeiner, S., Noebels, J.L., and Mucke, L. (2007). Aberrant Excitatory Neuronal Activity and Compensatory Remodeling of Inhibitory Hippocampal Circuits in Mouse Models of Alzheimer’s Disease. Neuron 55, 697–711.

Paxinos, G., and Keith B. J. Franklin, M. (2007). The Mouse Brain in Stereotaxic Coordinates. Elsevier Science.

Percie Du Sert, N., Ahluwalia, A., Alam, S., Avey, M.T., Baker, M., Browne, W.J., Clark, A., Cuthill, I.C., Dirnagl, U., Emerson, M., Garner, P., Holgate, S.T., Howells, D.W., Hurst, V., Karp, N.A., Lazic, S.E., Lidster, K., Maccallum, C.J., Macleod, M., Pearl, E.J., Petersen, O.H., Rawle, F., Reynolds, P., Rooney, K., Sena, E.S., Silberberg, S.D., Steckler, T., and Wurbel, H. (2020). Reporting animal research: Explanation and elaboration for the ARRIVE guidelines 2.0. PLoS Biol 18, e3000411.

Preuss, C., Pandey, R., Piazza, E., Fine, A., Uyar, A., Perumal, T., Garceau, D., Kotredes, K.P., Williams, H., Mangravite, L.M., Lamb, B.T., Oblak, A.L., Howell, G.R., Sasner, M., Logsdon, B.A., Consortium, M.-A., and Carter, G.W. (2020). A novel systems biology approach to evaluate mouse models of late-onset Alzheimer’s disease. Mol Neurodegener 15, 67.

Sanchez, P.E., Zhu, L., Verret, L., Vossel, K.A., Orr, A.G., Cirrito, J.R., Devidze, N., Ho, K., Yu, G.-Q., Palop, J.J., and Mucke, L. (2012). Levetiracetam suppresses neuronal network dysfunction and reverses synaptic and cognitive deficits in an Alzheimer’s disease model. Proceedings of the National Academy of Sciences 109, E2895–E2903.

Schneider, F., Baldauf, K., Wetzel, W., and Reymann, K.G. (2014). Behavioral and EEG changes in male 5xFAD mice. Physiol Behav 135, 25–33.

Sensini, F., Inta, D., Palme, R., Brandwein, C., Pfeiffer, N., Riva, M.A., Gass, P., and Mallien, A.S. (2020). The impact of handling technique and handling frequency on laboratory mouse welfare is sex-specific. Scientific Reports 10.

Stafstrom, C.E. (2005). The role of the subiculum in epilepsy and epileptogenesis. Epilepsy Curr 5, 121–129.

Surges, R., Volynski, K.E., and Walker, M.C. (2008). Review: Is levetiracetam different from other antiepileptic drugs? Levetiracetam and its cellular mechanism of action in epilepsy revisited. Therapeutic Advances in Neurological Disorders 1, 13–24.

Team, R.C. (2023). “_R: A Language and Environment for Statistical Computing_.”. 4.3.0 ed. (Vienna, Austria: R Foundation for Statistical Computing).

Toniolo, S., Sen, A., and Husain, M. (2020). Modulation of Brain Hyperexcitability: Potential New Therapeutic Approaches in Alzheimer’s Disease. Int J Mol Sci 21.

Tran, T.T., Speck, C.L., Pisupati, A., Gallagher, M., and Bakker, A. (2017). Increased hippocampal activation in ApoE-4 carriers and non-carriers with amnestic mild cognitive impairment. Neuroimage Clin 13, 237–245.

Veronese, M., Moro, L., Arcolin, M., Dipasquale, O., Rizzo, G., Expert, P., Khan, W., Fisher, P.M., Svarer, C., Bertoldo, A., Howes, O., and Turkheimer, F.E. (2019). Covariance statistics and network analysis of brain PET imaging studies. Scientific Reports 9.

Vossel, K., Ranasinghe, K.G., Beagle, A.J., La, A., Ah Pook, K., Castro, M., Mizuiri, D., Honma, S.M., Venkateswaran, N., Koestler, M., Zhang, W., Mucke, L., Howell, M.J., Possin, K.L., Kramer, J.H., Boxer, A.L., Miller, B.L., Nagarajan, S.S., and Kirsch, H.E. (2021). Effect of Levetiracetam on Cognition in Patients With Alzheimer Disease With and Without Epileptiform Activity. JAMA Neurology 78, 1345.

Vossel, K.A., Ranasinghe, K.G., Beagle, A.J., Mizuiri, D., Honma, S.M., Dowling, A.F., Darwish, S.M., Van Berlo, V., Barnes, D.E., Mantle, M., Karydas, A.M., Coppola, G., Roberson, E.D., Miller, B.L., Garcia, P.A., Kirsch, H.E., Mucke, L., and Nagarajan, S.S. (2016). Incidence and impact of subclinical epileptiform activity in Alzheimer’s disease. Annals of Neurology 80, 858–870.

Wan, Y.-W., Al-Ouran, R., Mangleburg, C.G., Perumal, T.M., Lee, T.V., Allison, K., Swarup, V., Funk, C.C., Gaiteri, C., Allen, M., Wang, M., Neuner, S.M., Kaczorowski, C.C., Philip, V.M., Howell, G.R., Martini-Stoica, H., Zheng, H., Mei, H., Zhong, X., Kim, J.W., Dawson, V.L., Dawson, T.M., Pao, P.-C., Tsai, L.-H., Haure-Mirande, J.-V., Ehrlich, M.E., Chakrabarty, P., Levites, Y., Wang, X., Dammer, E.B., Srivastava, G., Mukherjee, S., Sieberts, S.K., Omberg, L., Dang, K.D., Eddy, J.A., Snyder, P., Chae, Y., Amberkar, S., Wei, W., Hide, W., Preuss, C., Ergun, A., Ebert, P.J., Airey, D.C., Mostafavi, S., Yu, L., Klein, H.-U., Carter, G.W., Collier, D.A., Golde, T.E., Levey, A.I., Bennett, D.A., Estrada, K., Townsend, T.M., Zhang, B., Schadt, E., De Jager, P.L., Price, N.D., Ertekin-Taner, N., Liu, Z., Shulman, J.M., Mangravite, L.M., and Logsdon, B.A. (2020). Meta-Analysis of the Alzheimer’s Disease Human Brain Transcriptome and Functional Dissection in Mouse Models. Cell Reports 32, 107908.

Wang, M., Beckmann, N.D., Roussos, P., Wang, E., Zhou, X., Wang, Q., Ming, C., Neff, R., Ma, W., Fullard, J.F., Hauberg, M.E., Bendl, J., Peters, M.A., Logsdon, B., Wang, P., Mahajan, M., Mangravite, L.M., Dammer, E.B., Duong, D.M., Lah, J.J., Seyfried, N.T., Levey, A.I., Buxbaum, J.D., Ehrlich, M., Gandy, S., Katsel, P., Haroutunian, V., Schadt, E., and Zhang, B. (2018). The Mount Sinai cohort of large-scale genomic, transcriptomic and proteomic data in Alzheimer’s disease. Scientific Data 5, 180185.

Xiang, X., Wind, K., Wiedemann, T., Blume, T., Shi, Y., Briel, N., Beyer, L., Biechele, G., Eckenweber, F., Zatcepin, A., Lammich, S., Ribicic, S., Tahirovic, S., Willem, M., Deussing, M., Palleis, C., Rauchmann, B.S., Gildehaus, F.J., Lindner, S., Spitz, C., Franzmeier, N., Baumann, K., Rominger, A., Bartenstein, P., Ziegler, S., Drzezga, A., Respondek, G., Buerger, K., Perneczky, R., Levin, J., Hoglinger, G.U., Herms, J., Haass, C., and Brendel, M. (2021). Microglial activation states drive glucose uptake and FDG-PET alterations in neurodegenerative diseases. Sci Transl Med 13, eabe5640.

