## Supplementary Figures for "Levetiracetam Modulates Brain Metabolic Networks and Transcriptomic Signatures in the 5XFAD Mouse Model of Alzheimer’s disease"

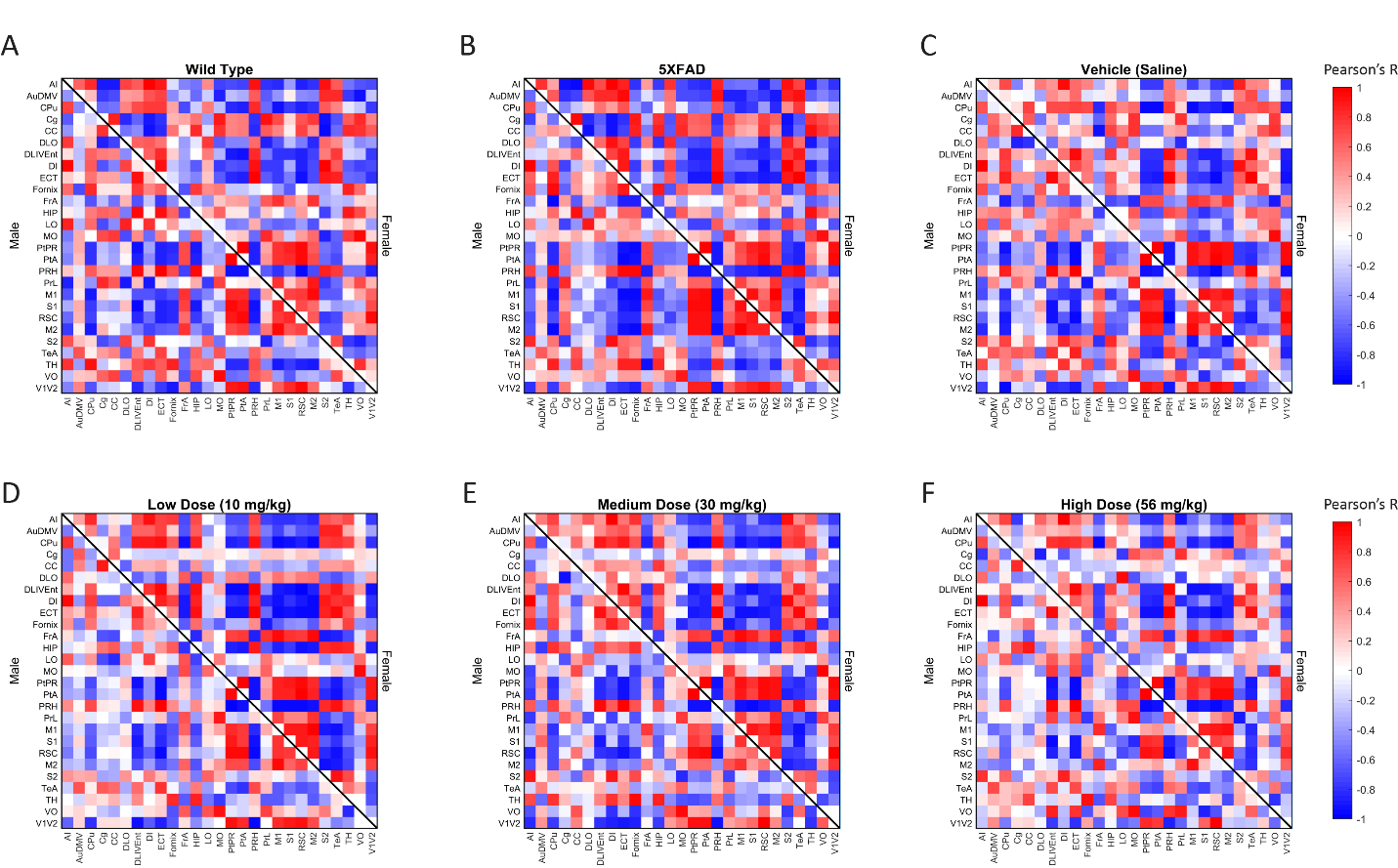

**Supplementary Figure 1**: Unthresholded metabolic covariance matrices sorted alphabetically by region.

**Supplementary Figure 2**: Pathway prediction legend for GO term visualization (Figure 7B-D).

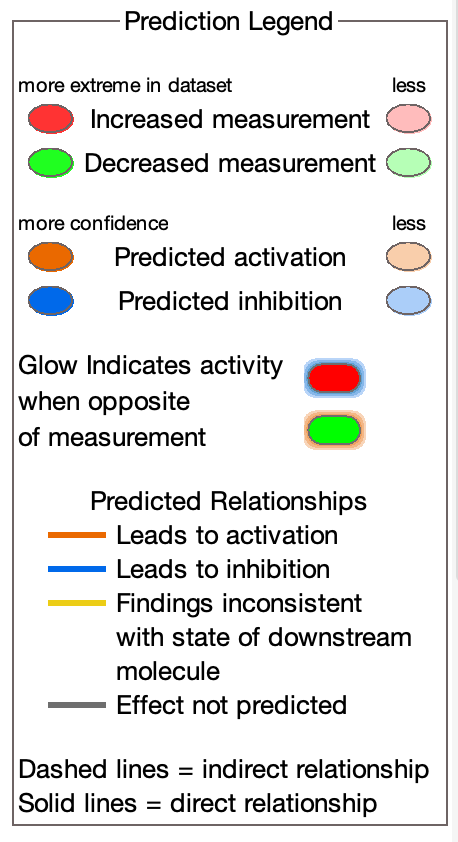

**Supplementary Table 1**: Full names of network nodes (region of interest) as defined in the Paxinos and Franklin Atlas. Left and right regions were averaged to yield a network of 27 regions normalized by the cerebellum..

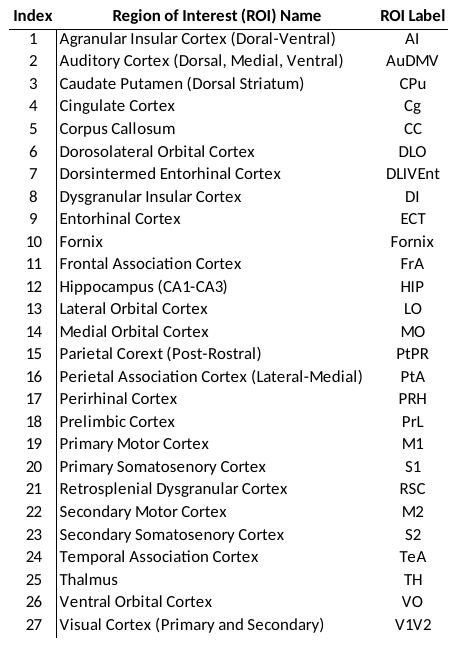
